# Kinesin-5 forms a stable bipolar spindle in a fast, irreversible snap

**DOI:** 10.1101/452821

**Authors:** Allen Leary, Elena Nazarova, Shannon Sim, Kristy Shulist, Paul Francois, Jackie Vogel

## Abstract

**GRAPHICAL ABSTRACT:** 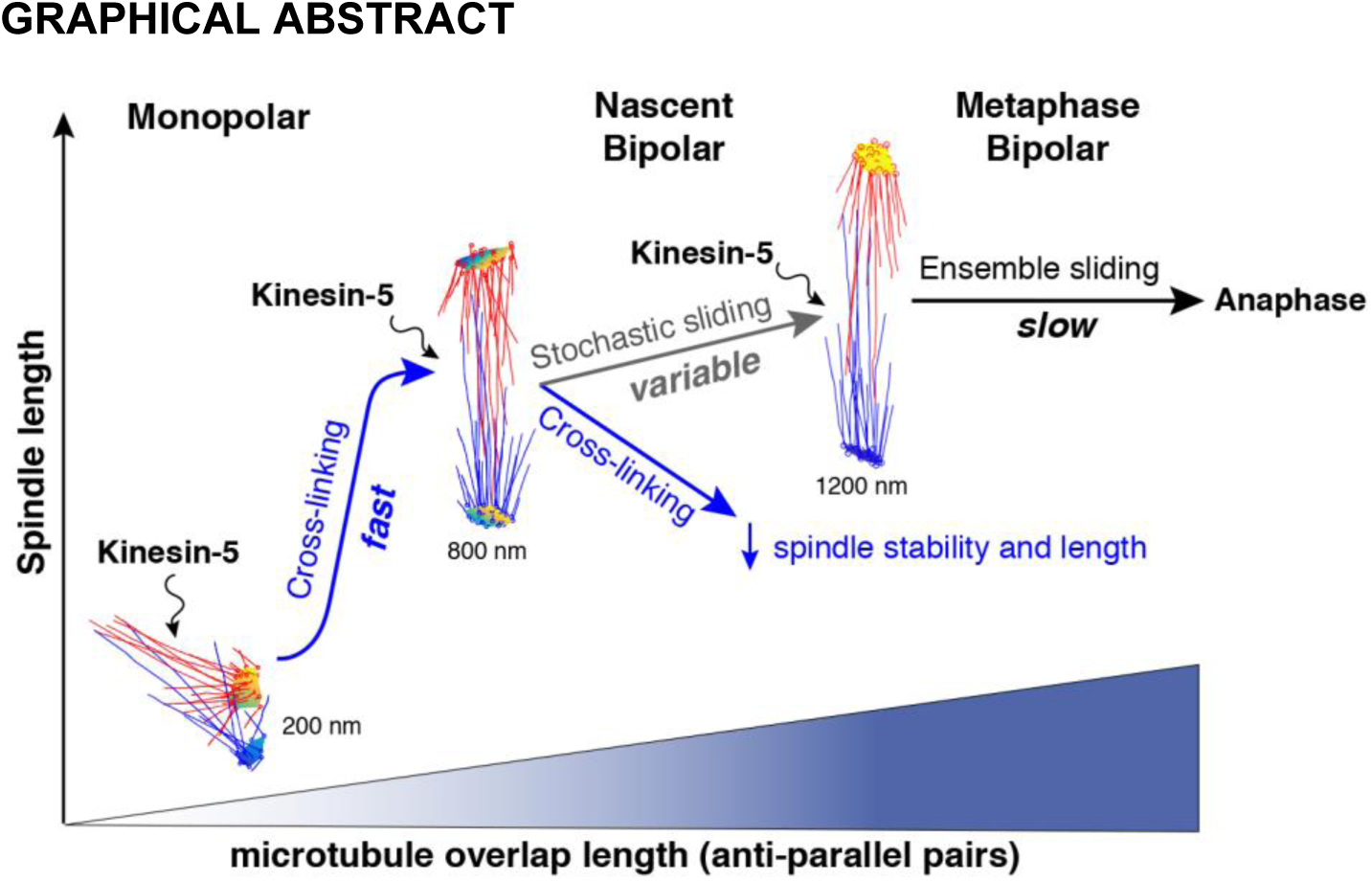

Separation of duplicated spindle poles is the first step in forming the mitotic spindle. Kinesin-5 crosslinks and slides anti-parallel microtubules, but it is unclear how these two activities contribute to the first steps in spindle formation. In this study we report that in monopolar spindles, the duplicated spindle poles snap apart in a fast and irreversible step that produces a nascent bipolar spindle. Using mutations in Kinesin-5 that inhibit microtubule sliding, we show crosslinking alone drives the fast, irreversible pole separation. Electron tomography revealed microtubule pairs in monopolar spindles have short overlaps that intersect at high angles and are unsuited for ensemble Kinesin-5 sliding. However, maximal extension of a subset of microtubule pairs approaches the length of nascent bipolar spindles and is consistent with a Kinesin-5 crosslinking driven transition. Finally, stochastic microtubule sliding by Kinesin-5 stabilizes the nascent spindle and sets a stereotyped equilibrium length.

## INTRODUCTION

Accurate segregation of chromosomes into daughter cells requires the formation of a bipolar spindle. The first step in forming a bipolar spindle is the transition from a monopolar state, where the duplicated spindle poles are unseparated, to the bipolar state. The bipolar spindle has a stereotyped design composed of two spindle poles on which microtubules (MTs) are assembled. MTs from opposing poles crosslink to form a set of stabilizing inter-polar MTs (ipMTs) while other MTs probe for, and attach to, the centromeres of sister chromatids. Kinesin-5 and Ase1 crosslink ipMTs, and mutations in these proteins cause spindle collapse (Hagan and Yanagida, 1992; Heck et al., 1993; Kapoor et al., 2000; Khmelinskii et al., 2007; Saunders and Hoyt, 1992). Two related and unresolved questions are how the organization of MTs in the monopolar state support the production of the forces required to separate the spindle poles, and why the nascent bipolar spindle does not collapse.

The formation of a bipolar spindle depends on the conserved Kinesin-5 family of plus end-directed motor proteins. Kinesin-5 is organized as a homo-tetramer and can both crosslink anti-parallel MTs as well as slide them apart (Kashina et al., 1996b). Studies to date have focused on Kinesin-5’s role in the maintenance of the bipolar spindle by virtue of its pushing forces which oppose the contractile forces enacted on the spindle by minus-end directed motors Kinesin-14 and dynein. Budding yeast has two Kinesin-5 orthologs: Cin8 and Kip1 (Hoyt et al., 1992; Straight et al., 1998b). Cin8 and Kip1 have been reported as being functionally redundant, however Cin8 is thought to have the greater role in forming a bipolar spindle as the Cin8 null mutant (cin8Δ), but not the Kip1 null mutant (kip1Δ), delays spindle formation, exhibits spindle instability and triggers activation of the spindle assembly checkpoint (Straight et al., 1998a). Mechanisms for bipolar spindle maintenance do not explain the role of transition from a monopolar state with duplicated spindle poles to a stable bipolar spindle. How Kinesin-5 drives this monopolar to bipolar transition through the MT architecture of the monopolar state remains unclear.

In this study we employ the simplicity of the budding yeast mitotic spindle to investigate the properties of the monopolar to bipolar transition and the contributions of both Kinesin-5 crosslinking and sliding modalities. We used a Cre-Lox recombinase-based fluorophore-switching system to spectrally separate spindle poles below the diffraction limit. Using simultaneous two-channel confocal microscopy, we track both spindle poles as individual point like objects before, during and after the monopolar to bipolar transition. Pole separation traces capture dynamic information for the transition and the dynamics of the newly formed spindle, allowing us to measure for the first time the velocity of the poles as they separate as well as the elongation rate and the stability of nascent bipolar spindles. The transition is fast and irreversible, with an average pole separation velocity of 17±4 nm sec^−1^, making it one of the fastest steps in spindle assembly. We report that mutations inhibiting Kinesin-5 MT sliding do not significantly change pole separation velocities. Using 3D electron tomography, we show that the MT structure of the initial monopolar state is comprised of short, high angle MT pairings that are unsuited to ensemble Kinesin-5 sliding. Nevertheless, this highly coupled system supports the maximal extension of crosslinked MTs required to achieve the bipolar state as measured for WT cells. We also find that the length and stability of the nascent bipolar spindle is dependent on Kinesin-5 MT sliding. We propose that while Kinesin-5 sliding does not specifically contribute to the monopolar to bipolar transition, both Kinesin-5 sliding and crosslinking are required to form a stereotyped, stable bipolar spindle.

## RESULTS

### The transition from monopolar to bipolar spindle is a fast, irreversible snap

Previous studies in both fission yeast (Rincon et al., 2017) and budding yeast (Crasta et al., 2006) demonstrated that Cin8 MT sliding is dispensable for forming a bipolar spindle. However, the velocity, irreversibility and efficiency of the transition in the presence or absence of Kinesin-5 sliding has not yet been measured. We defined spindle formation as a transition that links the monopolar spindle state, where duplicated spindle poles are unseparated, to the bipolar spindle state (Figure1A). In the monopolar state, the spindle poles are ~ 200 nm apart. The length distribution of bipolar pre-anaphase spindles in WT cells is 800 - 2100 nm (Nazarova et al., 2013; Shulist et al., 2017). To capture the transition, we employed a fluorophore switching technique (Hotz et al., 2012; Verzijlbergen et al., 2010) to spectrally separate and track the displacements of the pre-existing (old) pole from the newly assembled (new) pole. Spc42, a major component of the central plaque of the budding yeast spindle pole (Bullitt et al., 1997) was tagged with a Cre-Lox recombinase system that recombines out an in-frame mCherry tag and leaves an in-frame EGFP tag (Figure S1). As a result, monopolar spindles have an old pole labelled with Spc42-mCherry (Spc42-MCh) and a new pole labelled with Spc42-eGFP, each tracked as single point-like objects during spindle formation. This method provided a spatio-temporal resolution of 100 nm (in x,y and z) and 20 seconds and resolves the displacements of the unseparated spindle poles in the monopolar spindle state as well as during and after the transition (Figure 1B).

**Figure 1.**
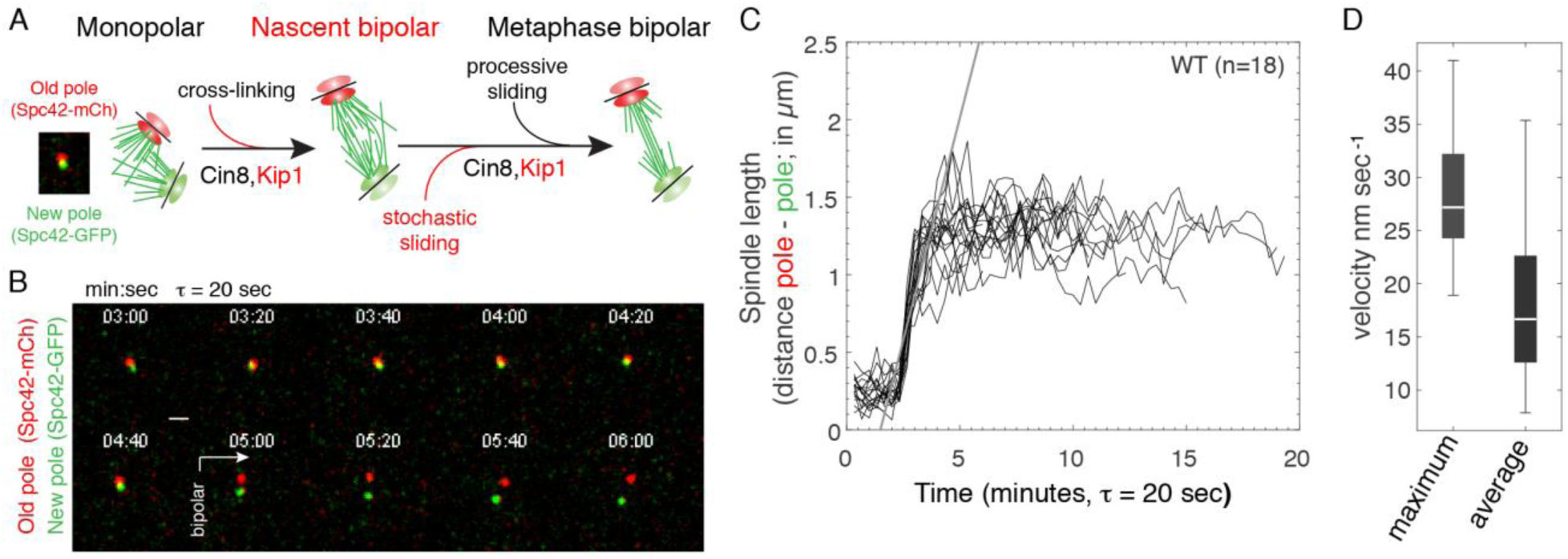
Characterizing the monopolar to bipolar spindle transition in living cells. (A) Working model for the monopolar to bipolar transition and the properties of nascent bipolar spindles. Contributions to early steps in spindle formation identified in this study are indicated in red. Our tracking method identifies each pole, as a single point-like object below the diffraction limit. Spectral separation is achieved by the old pole labelled with Spc42-mCh and the new pole with Spc42-GFP. The method used for fluorophore switching of Spc42 is described in Figure S1. (B) Representative lapse images of a wild type cell where Spc42-mCh labels the old spindle pole and Spc42-GFP labels the new spindle pole. Scale bar = 1 µm, time step τ = 20 seconds (C) Aligned traces of WT spindle length dynamics before, during and after the monopolar to bipolar spindle transition, with the slope for the maximum velocity of pole separation. Time step τ = 20 seconds. (D) Distributions for the maximum and average velocities of pole separation.

Using simultaneous dual camera imaging, we captured dynamic profiles that encompass the monopolar state, the transition and the new bipolar state. These dynamic profiles of asynchronous cells going through the transition can be aligned to give a stereotyped profile with the slope of the maximum velocity during the transition (Figure 1C). We defined the velocity of bipolar spindle formation as the velocity of pole separation between the points determined as the start and end point of the transition, defined by the local maxima/minima of the acceleration curves of the pole dynamic profiles. Surprisingly, the monopolar to bipolar transition is very fast in WT cells (*n*=18), with a maximal velocity of 28.7 ± 6.4 nm sec ^−1^ and average velocity of 17.1 ± 3.7 nm sec^−1^ (Figure 1D). The average velocity of pole separation during the monopolar to bipolar transition is similar to the previously reported velocity of ensemble Cin8 anti-parallel MT sliding (17 nm sec^−1^) measured in vitro (Adina et al., 2011), and is faster than any other transition in spindle assembly as well as fast anaphase (10-12 nm sec^−1^)(Straight et al., 1998b). The transition is irreversible, as we did not observe any instance of a nascent bipolar spindle collapsing to the monopolar state, or observe post-transition lengths less than 600 nm, well above the 200 nm pole to pole distance of monopolar spindles.

### Separation of the spindle poles is driven by Kinesin-5 MT crosslinking

Kinesin-5 motor activity relies on its ability to stably associate with MTs (Gheber et al., 1999). However, mutations that reduce the sliding activity of Cin8 while conserving its crosslinking activity have been identified and characterized in vitro, allowing us to investigate the individual contribution of each modality to the formation of a stable bipolar spindle.

The two budding yeast Kinesin-5 orthologs are similar in structure (Figure 2A), both being homotetrameric motors (Hildebrandt et al., 2006) capable of both crosslinking and +end directed sliding of antiparallel MTs (Fridman et al., 2013). To characterize the contribution of Kinesin-5 MT sliding to the monopolar to bipolar transition, we used two conditions to isolate sliding from crosslinking. First, we isolated Cin8’s crosslinking modality from its sliding activity using a Cin8-R196K point mutation (Cin8-RK) in the α2 helix of the motor domain (Figure 2A). The R196K mutation was previously shown to severely reduce motor activity in motility assays in vitro (Gheber et al., 1999). Second, we combined the Cin8-RK mutation with a null mutation in Kip1 (kip1Δ), resulting in a Cin8-RK kip1Δ double mutant with severely diminished Kinesin-5 sliding activity. We then measured the monopolar to bipolar transition in both conditions.

**Figure 2.**
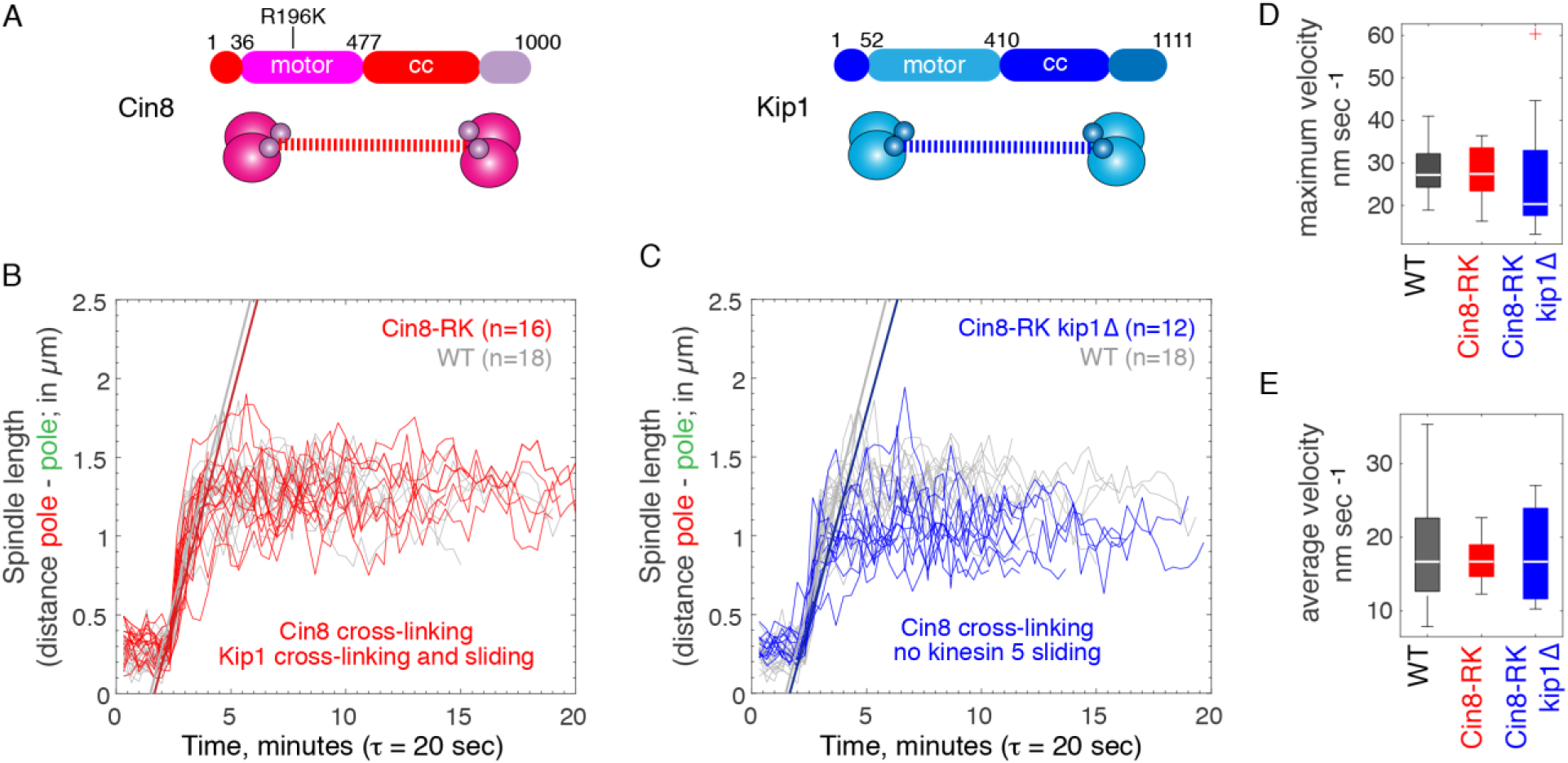
The monopolar to bipolar spindle transition is driven by Kinesin-5 cross-linking. (A) Cartoon depicting domains of Kinesin-5 orthologs Cin8 and Kip1. (B) Plots of spindle length dynamics during bipolar spindle formation in WT cells (—, *n*=18) and Cin8-RK mutant cells (—, *n*=16). (C) Plots of spindle length dynamics during bipolar spindle formation for both WT (—, *n*=18) and Cin8-RK kip1Δ cells (—, *n*=12).

The aligned traces of instantaneous length for the monopolar state, the transition and bipolar state of spindles in WT cells and the Cin8-RK mutant present remarkably similar profiles (Figure 2B). We found that the monopolar to bipolar transition in the Cin8-RK mutant (*n*=16) is fast, with a maximum velocity of 27.8 ± 5.9 nm sec ^−1^ (Figure 2D) and mean transition velocity of 16.9± 2.8 nm sec^−1^ (Figure 2E) that is not significantly different than either the maximum or mean velocity for WT cells (Figure 1D). The monopolar to bipolar transition is also irreversible in Cin8-RK cells. Our results show that fast, irreversible formation of a bipolar spindle formation does not depend on anti-parallel MT sliding by Cin8.

The two Kinesin-5 orthologs, Cin8 and Kip1, have overlapping roles in separating the spindle pole bodies apart, with Cin8 having the greater contribution (Hoyt et al., 1992; Roof et al., 1992). We reasoned Kip1 could contribute MT sliding activity in the Cin8-RK mutant. We found that as in WT cells and the Cin8-RK mutant, the monopolar to bipolar transition is fast and irreversible in the Cin8-RK kip1Δ mutant where all Kinesin-5 sliding was abolished (Figure 2C). As well, the maximum velocity and mean velocity of pole separation during the monopolar to bipolar transition in the Cin8-RK kip1Δ mutant is again remarkably similar to that of WT spindles; 26.7±13.4 nm sec ^−1^ and 17.8±6.5 nm sec^−1^ (*n*=12; Figure 2D and E). Together, these results show that fast, irreversible formation of a bipolar spindle does not require a significant anti-parallel MT sliding contribution from either Cin8 or Kip1.

### Crosslinked MTs in monopolar spindles extend to form a nascent bipolar spindle

The study of MT number and pairing using 3D electron tomography has provided many insights into structural changes that occur during spindle assembly and the metaphase to anaphase transition (Nazarova et al., 2013; O’Toole et al., 1999) but the majority of studies focus on metaphase or anaphase spindles. The MT organization required to initiate the separation of duplicated but unseparated spindle poles (a monopolar spindle) remains unclear and can provide insight into the evolution of spindle elongation and stability during the monopolar to bipolar transition. The nm scale resolution of MT lengths and interactions provided by 3D electron tomography allows us to correlate the system of MTs in the monopolar spindle and the dynamics we have measured for the monopolar to bipolar transition in living cells.

To enrich for wild type cells with monopolar spindles, we used elutriation to prepare samples for 3D electron tomography and were able to reconstruct four monopolar models (duplicated but unseparated poles with interacting MTs). As well, we identified a single 800 nm bipolar spindle, which represents an intermediate stage in metaphase (Nazarova et al., 2013; Winey et al., 1995). The length of this short bipolar spindle lies between the monopolar state (200 nm) and nascent bipolar state (~1300 nm), with 5% of bipolar spindles with a length of less than 1000 nm after the monopolar to bipolar transition (n=34 of 680 nascent bipolar spindle time points). While rare, the 800 nm spindle model provides important insight into the organization of MTs in nascent bipolar spindles.

We extracted information for key parameters that describe the MT architecture of the monopolar state and the transition to the nascent bipolar state; MT number, lengths, MT overlap pairings (length and angles; Figures S2 and S3 and videos S2-6). A representative model for a monopolar spindle is shown in Figure 3A and a short (800 nm) bipolar spindle is shown in Figure 3B. MTs emanating from the same spindle pole are coloured either blue or red. Overlap regions are shown in white, defined as the contours of MTs from opposing poles separated by ≤ 45 nm over ≥10 nm.

**Figure 3.**
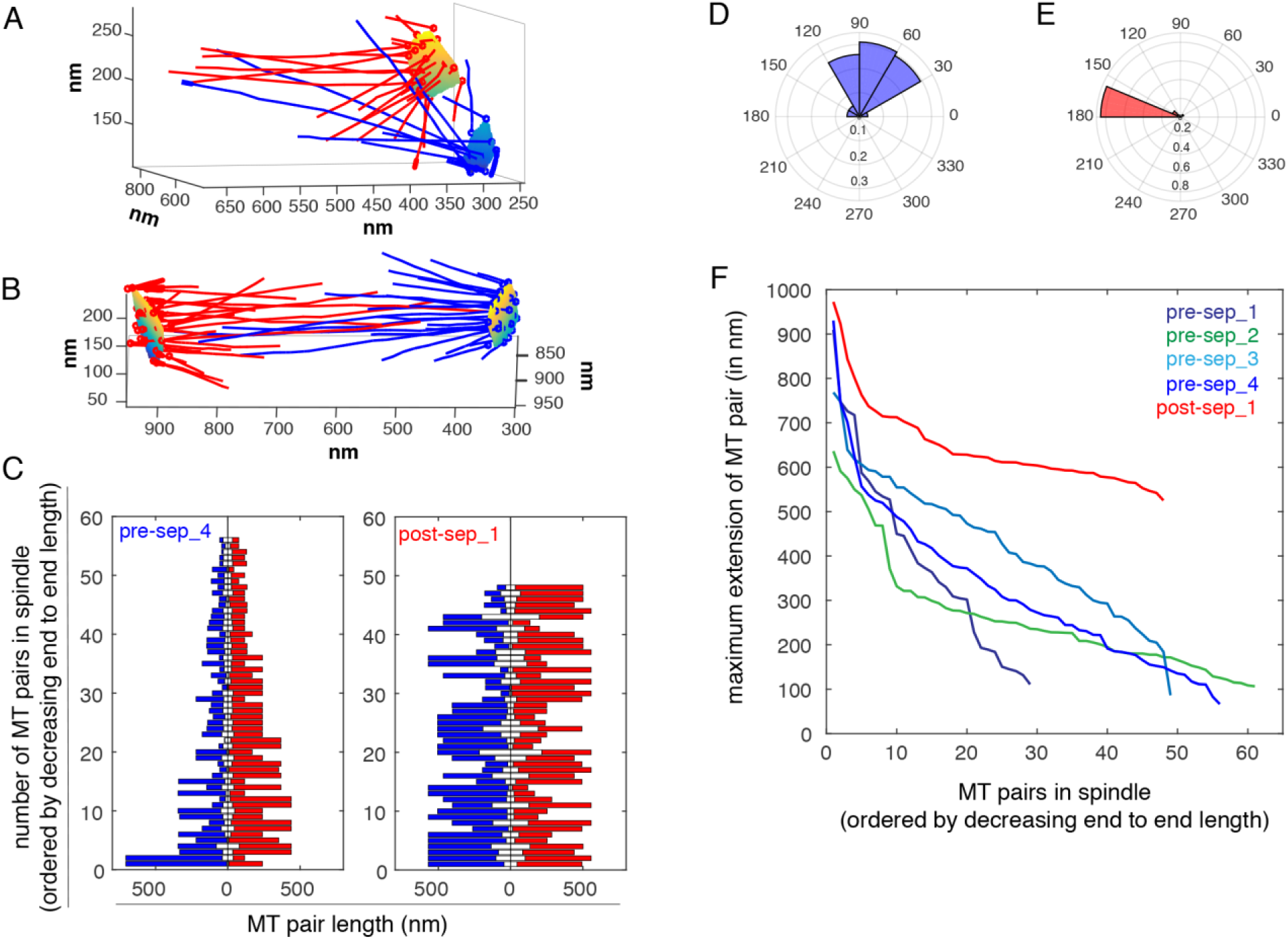
Microtubule architecture of monopolar and short bipolar spindles give insight into Kinesin-5 contributions. (A) Tomographic reconstruction of a monopolar spindle. (B) Tomographic reconstruction of a short (800 nm) bipolar spindle. For A and B, MTs of each pole are coloured blue or red. (C) Overlap profile of MT pairs. MTs pairs are spaced <45 nm with continuous overlap >10 nm. Blue and red indicate MTs from opposite poles, with the overlap region in white. Overlaps are sorted in descending order of total length for a monopolar spindle (left) and bipolar spindle (right). (D and E) Angles of MT pairs presented in a radar plot for the monopolar spindles and bipolar spindle shown in (A) and (B) respectively. (F) Maximum extension of the MT pairs as defined as the sum of their lengths minus the overlap length are plotted for 4 monopolar spindles and one 800 nm bipolar spindle. See also Figures S2 and S3, Table S1 and videos 2-6.

The four tomographic models of monopolar spindles possess between 29 to 61 distinct MT pairs (48.8 ± 14 MT pairs) with overlap lengths ranging from 11 nm to 159 nm (mean overlap 57.7 ± 23.7 nm). The 800 nm bipolar spindle has 48 distinct MT overlap pairs with lengths ranging from 13 nm to 434 nm (mean overlap 124.1 ± 104 nm). This contrasts with the overlap length of 400 nm that is typical of the ipMTs of metaphase spindles (Nazarova et al., 2013; Winey et al., 1995). As well, the number of intersecting MT pairs is much greater than the 6-8 ipMTs found in metaphase and anaphase spindles (Nazarova et al., 2013). This analysis reveals that monopolar spindles lack ipMTs and that the length of overlap between paired MTs in nascent bipolar spindles is 3-fold less than the overlap length of ipMTs in metaphase spindles. The overlap profiles of the electron tomography models are shown in Figure 3C (left unseparated and right early bipolar) with blue and red representing the lengths of each pair of MT from opposing spindle poles and in white the length of the overlap. Parameters for the overlap analysis are summarised in Table S1.

We also characterized the angles formed by the MT overlaps in the monopolar and 800 nm bipolar spindles as their organization is expected to give insight into crosslinking and/or sliding that can be supported by such an architecture. The MT pairs in monopolar spindles present a large range of overlap angles ranging from 30° to 160° but the majority (82%) of pairing angles are between 45° and 120° (Figure 3D). In the 800 nm bipolar spindle, the range of angles of MT pairs is more restricted than in the monopolar state, with 81% of MT pairs with angles ranging from 160° to 180° (with 180° being anti-parallel MTs; Figure 3E). Representative radar plots of angles formed by MT pairs of the tomogram models in Fig.3A and B are shown in Figure 3D and E. The results of our overlap angle analysis are summarised in Table S1.

Finally, we projected the maximal extension length (MEL) of the MT pairs extracted from the electron tomogram models. The MEL is the length obtained by summing the lengths of the two MTs that form the pair and subtracting the pairs measured overlap length. The MEL provides an upper bound for the length that the pair extends to, as it disregards where in the MT contour length and the angle the pairing occurs. We plotted the MEL for each MT pair for the four monopolar and 800 nm bipolar tomography models in Figure 3F. The MELs of monopolar spindles range between 600 nm and 900 nm but the vast majority of the MT pairs have a MELs below 500 nm. In contrast, the majority of MELs of the 800 nm bipolar spindle are >600 nm, with a minor set of MELs of 1000 nm. These long, paired MTs may be compressed. These results illustrate how MTs may be rearranged as a result of bipolar spindle formation. We assume MT crosslinking persists through the monopolar to bipolar spindle formation and that Kinesin-5 sliding does not significantly change the overlap length during the transition. With these two assumptions, we use the MELs (Figure 3F) to estimate the upper bound for the length of a bipolar spindle without the contribution of MT sliding. We approximate an upper bound of 600-900 nm, consistent with the overlap profile of the 800 nm bipolar model (600-1000 nm).

### The monopolar to bipolar transition is not delayed in Kinesin-5 mutants

We have shown that the rate and irreversibility of the monopolar to bipolar transition is not significantly altered in the absence of Kinesin-5 sliding. Moreover, our analysis of the MT architecture of monopolar spindles identified MT pairs with MELs that are consistent with the post-transition lengths of nascent bipolar spindles irrespective of Kinesin-5 sliding. Prior to bipolar spindle formation, Cdk1 restrains the proteolysis of Kinesin-5. This regulation ensures that sufficient Kinesin-5 is available to form MT pairs and drive formation of a bipolar spindle (Crasta et al., 2008). Kinesin-5 deficiencies have been found to delay the progression though stages of mitosis such as the duration of metaphase (Straight et al., 1998b) or anaphase (Movshovich et al., 2008). We considered the possibility that the monopolar to bipolar transition may equally be delayed in a Kinesin-5 sliding dependent fashion. If Kinesin-5 sliding contributes to the transition, then accumulation of a high concentration of Kinesin-5 motors with attenuated sliding activity may be sufficient to reach the force threshold required to separate the poles. A compensatory mechanism is expected to delay the onset of the transition in relation to the completion of spindle pole duplication. We therefore investigated if there was a significant cell cycle delay between START and the formation of a bipolar spindle in Cin8-RK and Cin8-RK kip1Δ cells relative to WT cells. START is the point of commitment to enter the cell division cycle, and can be used to benchmark the timing of spindle pole duplication in relation to pole separation, during which Kinesin-5 is expected to accumulate on the MTs of monopolar spindles (Crasta et al., 2006; Kraikivski et al., 2015; Skotheim et al., 2008). We used START as the beginning of the process of spindle formation in living cells.

The Whi5 transcriptional repressor is exported from the nucleus at START (Skotheim et al., 2008). We used Whi5-GFP nuclear export to measure the time from START (Skotheim et al., 2008) to the monopolar to bipolar transition. We used a timestep τ of 2 minutes, a timescale expected to capture the monopolar to bipolar transition in two consecutive timepoints. The transition was identified as the first time point where two distinct spindle poles labelled with Spc42-CFP were resolved (Figure 4A and video S7). We find the mean delay between START and bipolar spindle formation in WT cells is 71.1±23.5 mins (*n*=67; Figure 4B). The delay between START and bipolar spindle formation in Cin8-RK cells (64.4±23.5 mins; *n*=80) and Cin8-RK kip1Δ cells (68.9 ± 22.6 mins; *n*=88) was not significantly different than that of WT cells (Cin8-RK *p*=0.07; Cin8-RK kip1Δ *p*=0.70) (Figure 4B). The estimation statistics (Gardner and Altman, 1986) of the delay are plotted in Figure 4C, showing that the mean difference shift for both Cin8-RK and Cin8-RK kip1Δ lie within the 95% confidence interval. Our analysis demonstrates that loss of Kinesin-5 sliding activity does not delay the onset of the monopolar to bipolar transition after commitment to enter the cell cycle. The lack of cell cycle delay, together with indistinguishable monopolar to bipolar transition velocities for the Kinein-5 mutants relative to WT cells that is the outcome of a system of short MT overlaps, demonstrate MT crosslinking by Kinesin-5 is fully sufficient to undergo the monopolar to bipolar transition.

**Figure 4.**
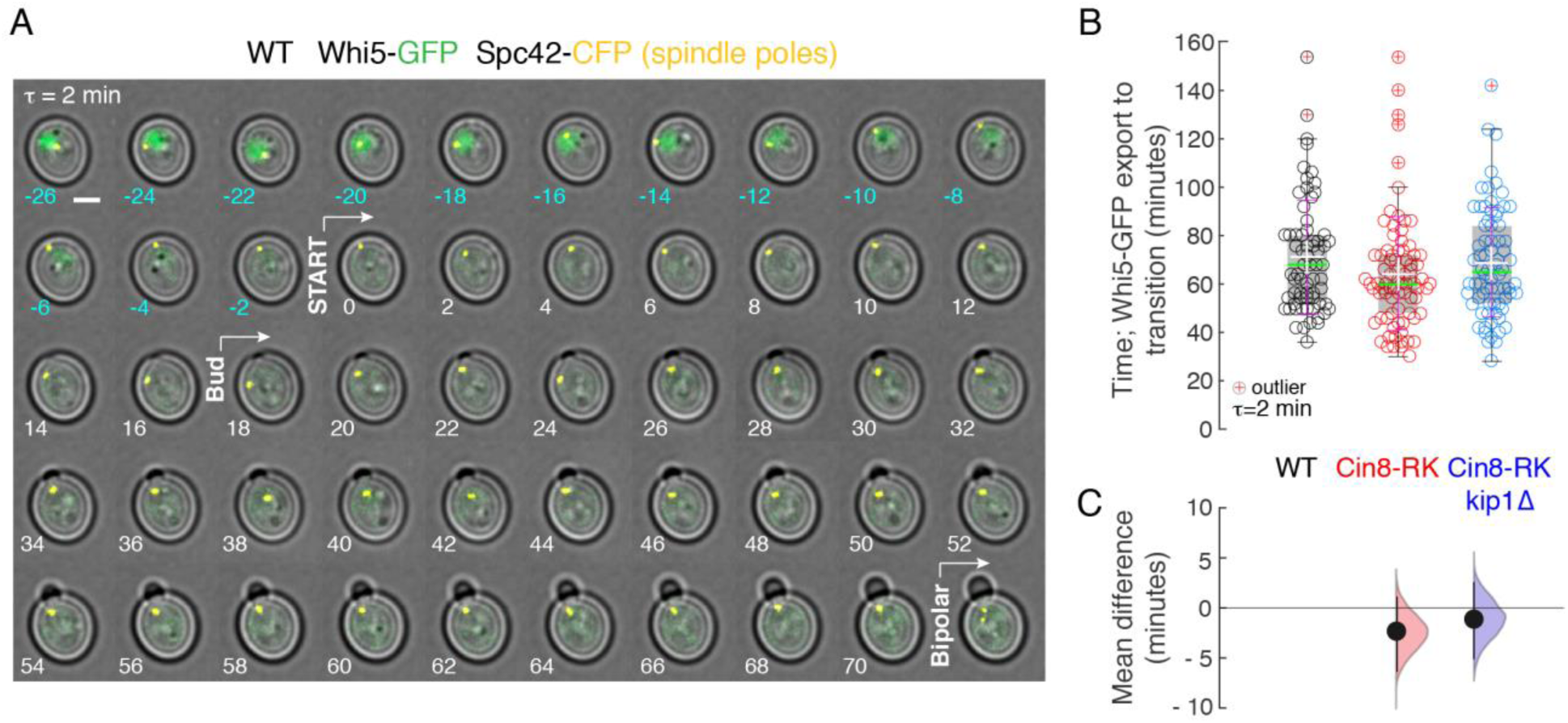
The monopolar to bipolar transition in Kinesin-5 mutants is not delayed relative to START. (A) Time lapse images of WT cells expressing Whi5-GFP showing timescale of nuclear export of Whi5. START (no Whi5-GFP signal in nucleus) and bud emergence are indicated on the timeseries. See also video S7. (B) Box plot with data points overlaid for the time delay between START and bipolar spindle formation in WT cells (○, *n*=67), Cin8-RK cells (○, *n*=80) and Cin8-RK Kip1Δ cells (○, *n*=74). (C) Estimation statistics plot of delays as visualized as a bootstrap 95% confidence interval of mean differences.

### Stochastic sliding of MTs by Kinesin-5 stabilizes the nascent bipolar spindle

The stability of the spindle is essential to form correct chromosome attachments and is the product of the concerted efforts of many highly regulated mitotic actors (van Heesbeen et al., 2014). Kinesin-5 has been shown to maintain spindle bipolarity by providing a plus ended sliding force (Kapitein et al., 2005; Kapoor et al., 2000; Kashina et al., 1996a) that counters the minus ended forces from Kinesin-14 (Goshima and Scholey, 2010). Previously, the stability of the bipolar spindle has been equated to simply the spindle lengths. Our data allow us to investigate for the first time the fluctuations in spindle length immediately after the transition to the nascent bipolar state. We therefore wanted to investigate the contribution of Kinesin-5 MT sliding and crosslinking to the stability and growth of the nascent bipolar spindle.

To understand how Kinesin-5 contributes to the dynamics of nascent spindles, we first determined if nascent spindles undergo net growth immediately following the transition to the bipolar state. We therefore extracted the mean velocity after the monopolar to bipolar transition (Figure 5A and Figure S4). The mean velocities obtained for the rate of elongation of nascent WT and Cin8-RK spindles are similar; respectively 0.21±0.429 and 0.27±0.393 nm sec^−1^, however we find that in Cin8-RK kip1Δ cells the majority of nascent spindles do not elongate (−0.143±0.307 nm sec^−1^). As a population, nascent spindles (WT, Cin8-RK and Cin8-RK kip1Δ) equally elongate or shrink, with a broad distribution of mean velocities illustrating a noisy system (−0.67 nm sec^−1^ to 1.2 nm sec^−1^; Figure 5A, bottom right).

**Figure 5.**
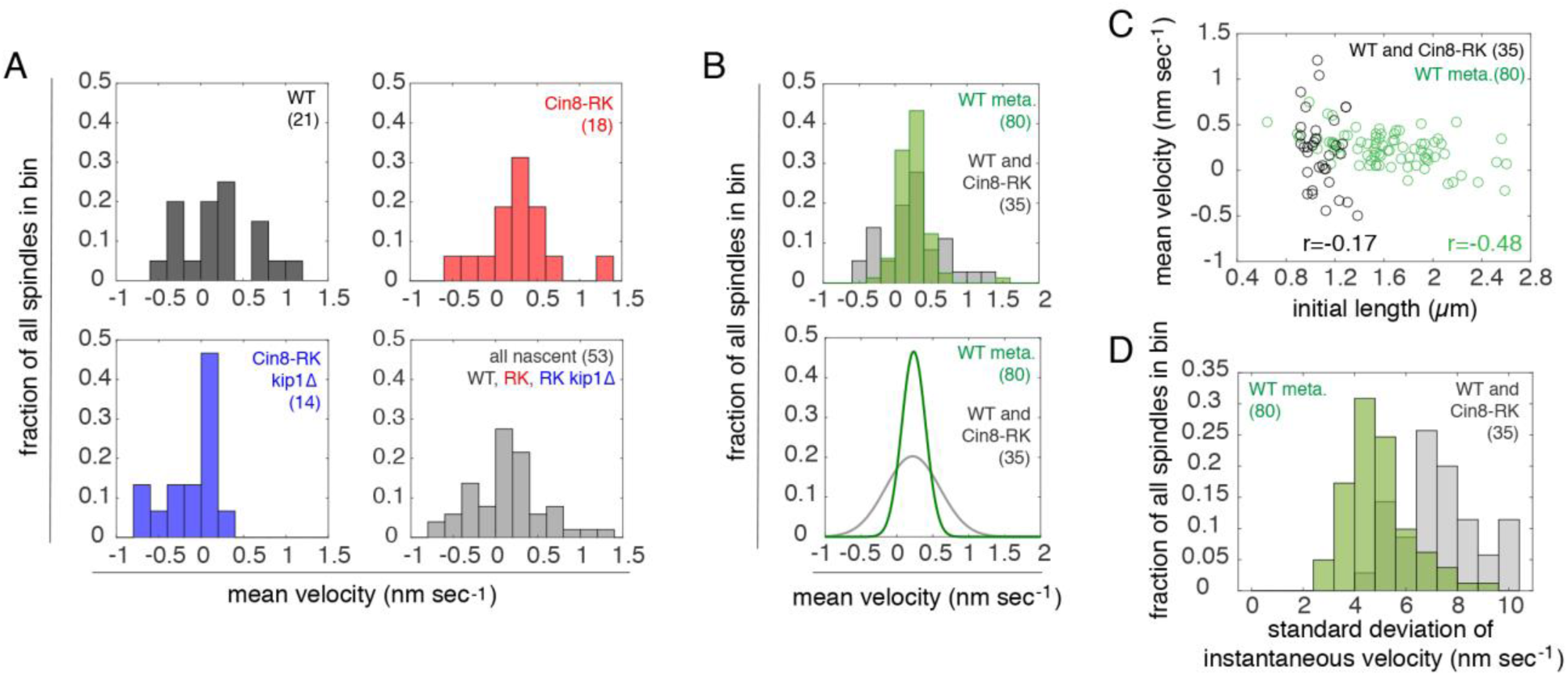
Stochastic elongation of nascent bipolar spindles. (A) Mean velocities of spindle elongation for nascent WT cells (top left; *n*=21), Cin8-RK cells (top right; *n*=18) and Cin8-RK Kip1Δ cells (bottom left; *n*=15). The combination of all spindle elongation rates (bottom right; *n*=53). (B) Mean velocity of spindle elongation for nascent WT and Cin8-RK spindles (*n*=39) and WT metaphase spindles (*n*=80). (C) Mean velocity of spindle elongation versus initial spindle lengths for WT nascent (**○**, *n*=21) and WT metaphase spindles (○, *n*=80) (D) Distribution of the standard deviation of instantaneous velocities of nascent and WT metaphase spindles. Also see Table S2.

Metaphase spindles grow prior to anaphase (Nazarova et al., 2013) and we therefore compared the growth of nascent bipolar spindles to that of a population of bipolar spindles with a length distribution that extends from those of nascent spindles to those of late metaphase. The growth parameters for nascent (WT, Cin8-RK and Cin8-RK kip1Δ) and metaphase spindles are provided in Table S2. The rate of growth for the metaphase spindle population (0.23±0.183 nm sec^−1^) was similar to the combined average for nascent spindles with Kinesin-5 sliding activity (WT and Cin8-RK; 0.24±0.4 09 nm sec^−1^). Interestingly, we found the variations in growth rate is halved in metaphase spindles, with nascent spindles exhibit a wider range of mean velocities (Figure 5B). The fraction of spindles undergoing net growth (Table S2) was increased in the metaphase population (0.925) relative to nascent WT (0.667) and Cin8-RK spindles (0.813). Our analysis shows that during metaphase, increasing spindle length is correlated with a transition to a stereotyped elongation regime.

Anaphase onset in WT budding yeast occurs at spindle lengths of 2.5-3 µ m (Straight et al., 1997). The nascent spindle has a mean length of 1.28 µm (Table S2) and therefore increases in length during metaphase. As spindle length is regulated (Goshima and Scholey, 2010) we asked if the length of the spindle influenced the velocity of spindle elongation. To compare nascent spindles to metaphase spindles, we correlated velocity to the initial length; either the length at the first timepoint after the transition for nascent spindles or the first timepoint of the time lapse for metaphase spindles. Interestingly, while the initial length of the nascent spindles has little correlation to the rate of elongation (Figure 5C, Pearson’s correlation coefficient r = −0.17), for the metaphase WT population the rate of elongation decreases as initial length increases (Nazarova et al., 2013) (Figure 5D, Pearson’s correlation coefficient r= −0.48). As well, nascent spindles exhibit much greater variation in instantaneous length relative to metaphase spindles (Figure 5D). Taken together, our analyses reveal that MT sliding by Kinesin-5 in nascent spindles is stochastic, but over time spindles switch to an ensemble elongation regime of net growth, where the rate decreases with the approach to anaphase.

Spindle elongation and stability are needed to bi-orient chromosomes in metaphase. Spindle instability or collapse are the cause of chromosome attachment defects and chromosome loss (Kapoor et al., 2000). We therefore next compared the stability of nascent spindles to that of spindles in the metaphase WT population to understand how stability evolves after the monopolar to bipolar transition. We measured spindle stability as a function of length fluctuations over the tracked time course, with the fluctuations normalized to mean spindle length (Nazarova et al., 2013; Shulist et al., 2017). Nascent spindles exhibit large length fluctuations with an amplitude that overlaps (*p* value > 0.4) with the fluctuations of short metaphase spindles with similar mean lengths (<1.499 µm; Figure 6A). However, as spindle length increases, length fluctuations are dampened (*p* value< 3·10^−8^ for metaphase WT spindles >1.499 µm; Figure 6A and C).

**Figure 6.**
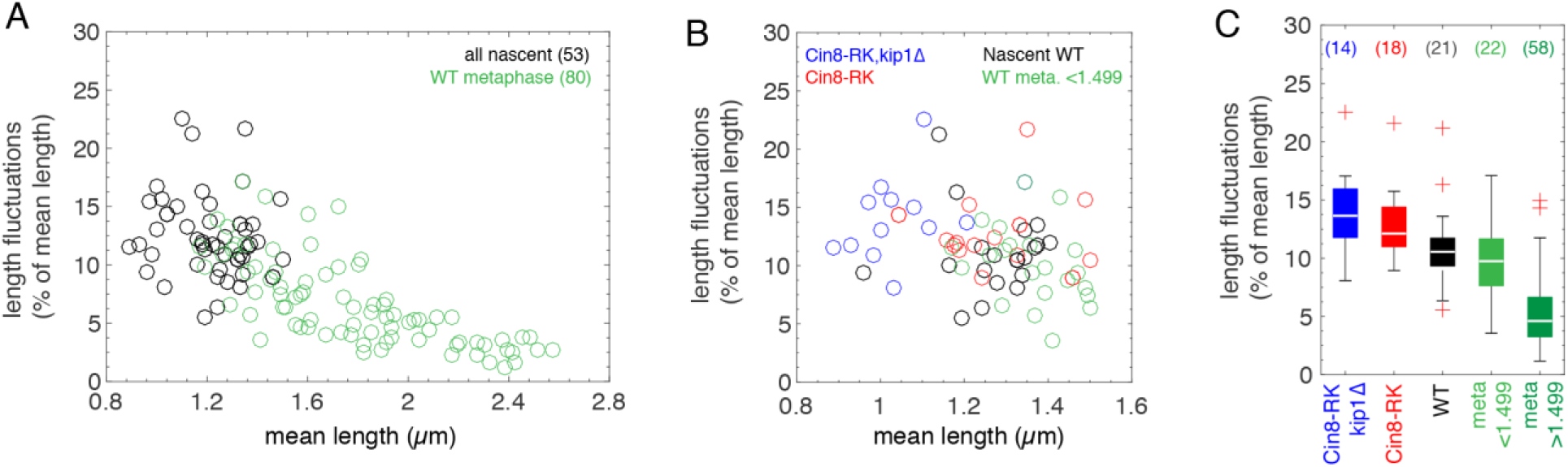
Kinesin-5 sliding dampens length fluctuations in nascent spindles. (A) Spindle length fluctuations in % of mean length versus mean spindle length for nascent (○; *n*=53) and WT metaphase (○; *n*=80) (B) Spindle length fluctuations of nascent Cin8-RK,kip1Δ (○;n=14), Cin8-RK (○; *n*=18), WT (○; *n*=21) and WT metaphase spindles of mean length < 1.499µm (○; *n*=22). (C) Box plot of spindle length fluctuations shown in (B). See also Figure S4.

We next asked if Kinesin-5 sliding is needed to stabilize nascent spindles. We measured normalized length fluctuations in nascent WT, Cin8-RK and Cin8-RK kip1Δ spindles and detected an increase in spindle fluctuations when Kinesin-5 sliding is deficient; with Cin8-RK nascent spindles exhibiting larger fluctuations than WT nascent spindles (*p* < 0.05) and Cin8-RK kip1Δ spindles exhibiting larger fluctuations than either Cin8-RK or WT nascent spindles (*p* = 0.001) (Figure 6B). Our results demonstrate that nascent spindles, which lack the ipMTs found in metaphase spindles, are stabilized by stochastic Kinesin-5 MT sliding.

### Kinesin-5 MT sliding sets the equilibrium length of nascent bipolar spindles

Cells lacking Cin8 have short metaphase spindles that are defective in chromosome attachment (Gardner et al., 2008; Straight et al., 1998b). These defects are the outcome of the dual role of Kinesin-5 in both spindle elongation as well as chromosome congression and kinetochore tension (Suzuki et al., 2018). We speculated that a combination of Kinesin-5 crosslinking and stochastic MT sliding ensures the nascent spindle attains a minimum “equilibrium” length that supports chromosome attachment. We computed the monopolar and nascent bipolar equilibrium length by binning the instantaneous spindle lengths for WT, Cin8-RK and Cin8-RK kip1Δ spindles (Figure 7A and B). A double gaussian fit is shown for each distribution, and the double gaussian fits for the WT, Cin8-RK and Cin8-RK kip1Δ distributions are shown together in Figure 7C. The equilibrium length of the monopolar state is defined by the peak and width of the gaussian fit for all cells, and is 240 – 300 nm across all three conditions, consistent with the ~200 nm distance between the duplicated spindle poles of monopolar spindles reported using electron tomography (Winey et al., 1995).

**Figure 7.**
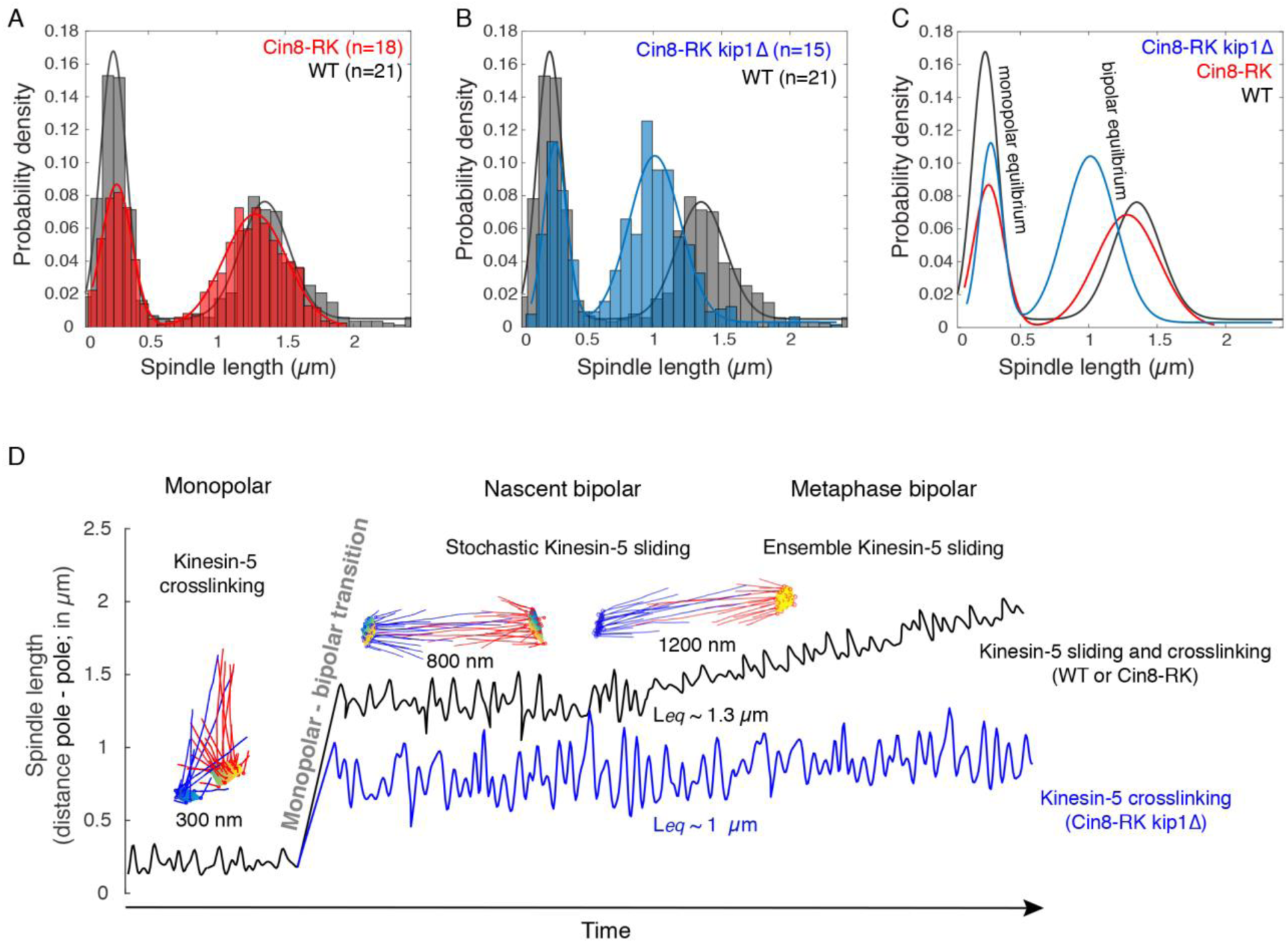
Kinesin-5 sliding sets the equilibrium length of new bipolar spindles. (A) Histograms of the relative probabilities of the spindle lengths during the monopolar to bipolar transition for WT cells (—) and Cin8-RK cells (—) with their associated double gaussian fits. (B) Histograms of the relative probabilities of the spindle lengths during the monopolar to bipolar transition for WT cells (—) and Cin8-RK Kip1Δ cells (—) with their associated double gaussian fits. (C) Overlay of the fits obtained from the probability distributions shown in (A) and (B). (D) A working model of Kinesin-5 contributions to the monopolar to bipolar transition and the stability and growth of a nascent bipolar spindle. Kinesin-5 crosslinking enacts a fast, irreversible and stereotyped transition from monopolar state to a bipolar spindle. Kinesin-5 sliding ensures that the equilibration length of the nascent spindle is sufficient to enact tension forces to correctly attach chromosomes and progress through mitosis. Kinesin-5 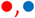; SPB, 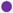; MTs, —.

The contribution of stochastic Kinesin-5 sliding to nascent bipolar spindle length was apparent when comparing the equilibrium length of WT (1.35±0.42 µ m) and Cin8-RK nascent spindles (1.28±0.54 µm) to that of Cin8-RK kip1Δ cells (1.01±0.54 µ m). Our results demonstrate that either Cin8 or Kip1 can slide MTs to set the equilibrium length of the nascent spindle. The equilibrium length of nascent bipolar spindles lacking a significant contribution of MT sliding by either Cin8 (Cin8-RK) or Kip1 (Cin8-RK kip1Δ) can be reconciled with the MELs of the longest MT pairs extracted from electron tomography models of monopolar spindles and the 800 nm bipolar spindle (Figure 3F). Short MT overlaps are expected to be stabilized by Kinesin-5 crosslinking but are unlikely to support processive Kinesin-5 sliding. Nevertheless, Kinesin-5 sliding does have a critical role in spindle formation, as it is required for the nascent bipolar spindle to attain a stable equilibrium length of 1.3 µm immediately following the transition (Figure 7D).

## DISCUSSION

### The monopolar to bipolar transition is a fast and irreversible process that produces a stable spindle

In this study, we demonstrate for the first time that the transition from a monopolar spindle to a bipolar spindle is faster than any other step in spindle assembly including anaphase. We report that once a bipolar spindle is formed, it remains bipolar and does not collapse back to a monopolar state, thus spindle formation is a two-state irreversible transition.

Using mutations in the two Kinesin-5 orthologs of budding yeast, we have dissected out the contributions of Kinesin-5 crosslinking and sliding modalities to the monopolar to bipolar transition. We found that the crosslinking modality of Kinesin-5 is sufficient for fast and irreversible formation of a bipolar spindle. We have shown that Kinesin-5 sliding nonetheless contributes to specifying the equilibration length of the nascent bipolar spindle as previously suggested (Straight et al., 1998a) and dampening length fluctuations. Dampening of spindle fluctuations occurs throughout metaphase spindle assembly up until anaphase. We equally show that the nascent bipolar spindles require Kinesin-5 to start undergoing net elongation towards the anaphase onset spindle lengths. The nature of this net elongation switches from a stochastic noisy regime in the nascent spindles to a stereotyped elongation regime as metaphase progresses only to slow down at the approach of anaphase onset lengths. Together our results highlight Kinesin-5 crosslinking as the driver of the formation of nascent bipolar spindles and sufficient to prevent their collapse. However, Kinesin-5 sliding is required to fully dampen the spindle’s fluctuations and allow it to undergo stereotyped elongation towards anaphase onset lengths.

### The microtubule architecture of monopolar spindles reveals the need for successive MT crosslinking and sliding activities of Kinesin-5 during spindle formation

Combining our study of the MT architecture and our characterization of the role of Kinesin-5 crosslinking in the monopolar to bipolar spindle transition has provided us with a comprehensive outlook on the sources of force generation for this transition. Our study has shown that the monopolar spindle contains many short and/or high angle (>30°) anti-parallel MT pairs. Cin8 appears to be particularly well suited to crosslink MTs across this diverse and challenging set of pairing architectures found in monopolar spindles (Shimamoto et al., 2015) thanks to its non-canonical MT binding sites (Bell et al., 2017). Cin8 also possesses a nonmotor MT binding site that is required for crosslinking as it increases its affinity to MTs independent of motor forces (Weinger et al., 2011). Cin8 has also been found to cluster near the poles of the monopolar spindles which puts it in a position to capture and crosslink MTs from the opposing pole (Shapira et al., 2017).

However, in the monopolar state the overlap length of MT pairs is often short (>100 nm) and not expected to support cooperative Kinesin-5 sliding responsible for the large anaphase spindle elongations (Roostalu et al., 2011; Shimamoto et al., 2015). These high angle crosslinks have been found to enact measurable deflection forces in vitro (Shimamoto et al., 2015). These forces, summed across many MT pairs as present in monopolar spindles could provide a considerable pushing force to separate the spindle poles. Individual Kinesin-5 motors on a crosslinked MT have been found to apply approximately 1.5 pN to the MT (Shimamoto et al., 2015). Our analysis of the MT organization within monopolar spindles shows up to 50 paired MTs, each of which will be crosslinked by Kinesin-5 motors. This could sum to forces in the hundreds of pNs range which is comparable to those during maximal pole separation (Cytrynbaum et al., 2003).

Kinesin-5 and the structural MAP Ase1 both stabilize bipolar spindles through MT crosslinking (Khmelinskii et al., 2009). In budding yeast, loss of Ase1 increases the frequency of spindle collapse in anaphase (Schuyler et al., 2003). Ase1 also functions to stabilize metaphase spindles. In both fission and budding yeasts, MT crosslinking appears to be sufficient for formation of a bipolar spindle. In fission yeast, Ase1 crosslinking alone supports bipolar spindle formation in the absence of both Kinesin-5 and the minus end directed motor Kinesin-14 (Rincon et al., 2017). In budding yeast, overexpression of Ase1 in G1 supports the formation of a bipolar spindle in the absence of both Cin8 and Kip1 (Crasta et al., 2006), however these cells are inviable. This suggests that MT crosslinking by Ase1 alone cannot support the formation of a functional bipolar spindle. Our finding that Kinesin-5 MT sliding is important for the nascent bipolar spindle to attain a stereotyped equilibrium length is consistent with Ase1 playing a minor and/or partially redundant role in spindle formation in budding yeast. Together with our findings, these studies reinforce the important role of MT crosslinking by Kinesin-5 in the formation of the eukaryotic bipolar spindle. Nevertheless, a detailed analysis that investigates Ase1’s contributions to the monopolar to bipolar transition and the stability of nascent spindles will be an important step towards fully understanding the mechanism for the formation of a stable bipolar spindle.

In both animal and fungal cells, the duplicated spindle poles of the monopolar spindle are tethered by a linker which must be cleaved in order to form a bipolar spindle. Cleavage of the linker structure therefore has a major role in regulating the progression of mitosis via the monopolar to bipolar transition. The full composition of the spindle pole linker is still unclear, but contains proteins conserved among eukaryotes including centrin and the coiled-coil protein Sfi1 (hSfi1 in human cells). Intriguingly the Sfi1 protein has been implicated in restricting one round of pole duplication per cell cycle via phospho-regulation of its C-terminus (Elserafy et al., 2014; Seybold et al., 2015). Sfi1C-termini dimerize to form a linker between the duplicated but unseparated spindle poles and these dimers are an important junction for regulating bridge cleavage and spindle pole separation. Sfi1 phospho-regulation has been implicated in delaying and altogether blocking Sfi1 dimer dissociation in budding yeast, implying a mechanism that regulates the cleavage of the half bridge structure might sense force.

The force acting on the half bridge could produce the tension that is required for Sfi1 dimer dissolution. The half bridge may store tension that is released when the threshold is met for monopolar to bipolar spindle transition. This threshold requires sufficient Kinesin-5 crosslinking to trigger the transition. The accumulation of forces on the linker up to a certain threshold in combination with cell cycle related phospho-regulation can provide a tension-sensing mechanism that produces the bimodal step-like transition we observe for the monopolar to bipolar transition.

### An irreversible transition to a stable nascent bipolar state is important for subsequent steps in mitosis

Kinesin-5 has diverse roles in mitosis, arising from its ability to crosslink and slide MTs. Its processive plus ended sliding of ipMTs has been seen as the main driver of spindle elongation and as a result, chromosome segregation (Stephens et al., 2013). But recent studies have revealed a plethora of different properties and roles for Kinesin-5, namely additional non-canonical MT binding sites (Bell et al., 2017; Weinger et al., 2011) in addition to binding sites in the motor domain required for motility (Adina et al., 2011; Edamatsu, 2014; Fridman et al., 2013; Roostalu et al., 2011). Kinesin-5 also localizes to the kinetochore-MT interface and acts as a length dependent depolymerase to facilitate the congression of chromosomes (Gardner et al., 2008), and recruits PP1 to kinetochores, contributing to maintaining Ndc80 attachment to MTs and tension generation (Suzuki et al., 2018). Our study reveals that an essential step in mitotic spindle assembly that had been previously overlooked as simply a result of Kinesin-5’s sliding modality in fact integrates both its sliding and crosslinking modality to produce a stable and elongating metaphase spindle (Fig 7D). In all cell types the length of the metaphase spindle is stereotyped to the volume of the cells (Goshima and Scholey, 2010). Maintaining this length is the result of the balanced interplay between many actors, and so disrupting any one actor such as MT dynamics, kinetochore attachments or MT associated proteins, in particular mitotic motor proteins such as Kinesin-5, can severely affect the faithful segregation of genetic material.

Stable spindle bipolarity is essential for correct chromosome attachment in mitosis (Tanaka, 2010; Tanenbaum and Medema, 2010) and meiosis (Segbert et al., 2003). The mechanisms by which cells sense and maintain correct MT-kinetochore attachments rely on both attachment and tension, but maintaining the bipolar state is the first and most important requirement (Musacchio and Salmon, 2007; Pinsky and Biggins, 2005). Correct chromosome attachment is a noisy and stochastic process sensitive to small defects in MT dynamic instabilities and the interactions between many protein complexes that govern the kinetochore-MT interface (Gay et al., 2012; Maiato et al., 2004). Spindle collapse is expected to severely impair mechanisms for stabilizing kinetochore-MT attachments emanating from opposing poles. In a collapsed spindle, the likelihood of syntelic and merotelic attachments increases due to the proximity of MTs emanating from both poles (Tanaka, 2010). When the mammalian Kinesin-5 Eg5 is inhibited by monastral, chromosome dispersion (a measure of defective chromosome congression) is maximal when metaphase spindle length is reduced by only 35% (Kapoor et al., 2000).

The threshold between pathological spindle instability leading to chromosome loss and “physiological” instability that is characteristic of nascent bipolar spindles is unclear. The intrinsic instability of the nascent bipolar spindle is not captured in the dynamic models used to understand the chromosome search and capture mechanism, which assume that the positions of the spindle poles are static when simulating MT dynamics, kinetochore-MT attachments and chromosome movements (Paul et al., 2009; Vasileva et al., 2017; Wollman et al., 2005). The stereotyped length of nascent bipolar spindles provides insight into the dual role of Kinesin-5 in setting the length threshold. Our findings reveal that coupling Kinesin-5 crosslinking and MT sliding during the earliest steps in spindle formation ensures a rapid transition to a stable bipolar state that supports the efficient search and capture of kinetochores as well as a sufficient length for tension to be applied to bioriented chromosomes.

## AUTHOR CONTRIBUTIONS

Conceptualization, J.V. and A.L.; Methodology, A.L. and E.N.; Formal Analysis, A.L. and S.S.; Investigation, A.L., E.N., S.S. and K.S.; Writing-Original Draft A.L. and J.V.; Writing-Review and Editing, A.L., J.V., S.S, K.S. and P.F.; Supervision, J.V. and P.F.; Funding Acquisition J.V. and P.F.

## ACKNOWLEDGEMENTS

The authors thank members of the Vogel lab, Stephanie Weber, Susanne Bechstedt and Gary Brouhard for discussions and their comments on the manuscript, the Integrated Quantitative Biology Initiative for access to the confocal microscope, the FEMR facility at McGill University for access to high pressure freezing equipment used to prepare cells for monopolar spindle EM tomography. Tomography was performed with Eileen O’Toole at the Boulder Electron Microscopy Services at the University of Colorado, Boulder. AL is supported by fellowships from the Cellular Dynamics of Macromolecular Complexes training program (Natural Sciences and Engineering Research Council of Canada), the Fonds Nature et technologies Quebec (FRQNT) and the Lorne Trottier Science accelerator award. This research was supported by a grant from the Canadian Institutes of Health Research (MOP-123335) to JV, a FQRNT team grant (G241608) to PF and JV, and an Innovation Fund infrastructure grant (33122) from the Canadian Foundation for Innovation to JV.

## DECLARATION OF INTERESTS

The authors declare no competing interests.

## LIST OF SUPPLEMENTAL MATERIALS

## 1. Supplemental methods

### Strain construction and cell culture conditions

All yeast strains are derivatives of BY4741. Mutant form of Cin8 (cin8-3) was generated in plasmid pSU19 using the USER cloning (New England Biolabs) (Bitinaite et al., 2007) PCR amplified and integrated into the CIN8 locus by homologous recombination. The resulting strain was crossed with a haploid strain which carries the Fluorophore switching cassette, loxP-mCherry-KanMx4-loxP-GFP His3-MX6 (Hotz et al., 2012). The haploids bearing CIN8-R196K in combination with the fluorophore-switching cassette were isolated by tetrad dissection. The conditional estradiol inducible Cre recombinase from pYB1109 (Hotz et al., 2012) was randomly integrated into the genome of EN15 and YV1085 by transformation resulting in EN1 and YV1099 strains, respectively. EN1, YV1099 and YB1037 strains were used for microscopy.

### Live cell microscopy

Yeast strains were grown to log phase (OD_600 ~0.1-0.6) in synthetic complete (SC) media supplemented with 2mM ascorbate at 25°C. All live cell imaging was undertaken at 25°C. For bipolar spindle formation and fluorophore switching technique Cre recombinase induction was used, β-Eestradiol (1μM) was added to SC media 120 min before imaging. Cells were prepared for imaging as previously reported(Nazarova et al., 2013; Shulist et al., 2017)

Bipolar spindle formation was imaged on a Quorum Diskovery platform installed on a Leica DMi8 inverted microscope. This system consists of a HCX PL APO 100x/1.47 OIL CORR TIRF objective (Leica), an DISKOVERY multi-modal imaging system (Spectral) with a multi-point confocal 50um pinhole spinning disk and dual iXon Ultra 512×512 EMCCD (Andor) cameras for simultaneous imaging, ASI three axis motorized stage controller, and MCL nano-view piezo stage, 488nm and 561nm solid state OPSL lasers linked to a Borealis beam conditioning unit. Image acquisition and microscope control was executed using MetaMorph (Molecular Devices). Images were acquired in stream mode, with each 6 µm z-stack composed of 30 z-slices of 200 nm, and an exposure time of 50 ms. Cells were imaged for 20 mins with a 20 s time step between stack acquisitions (60 time points).

To measure the timing of bipolar spindle relative to START, early log phase cells (OD600 0.1–0.3) were concentrated and spotted onto a pad of 1% agarose SC contained in a gene frame (ThermoScientific). Cells were incubated on the agar pad/slide for 10 mins at ambient temperature before imaging. Imaging data was taken on Olympus IX83 microscope using 100x oil objective lens (Olympus Plan Apo 100X NA 1.40 oil). Images were captured using a Hamamatsu Orca-Flash 4.0 sCMOS camera. Z-stacks were done using a NanoScanZ piezo by Prior Scientific. Excitation was undertaken using an X-Cite 120 LED lamp using the ET–ECFP/EYFP/mCherry filter set (Chroma). Olympus CellSens 2.1 imaging software was used to control the microscope, illumination and acquisition. Images were collected as 5 µm Z-stacks (10 slices of 0.5 µ m) with exposure time of 100 ms. Cells were imaged for 4 hours with a 2 min τ between stack acquisitions (120 timepoints). A 2 min τ was selected based on the ~ 90 sec timescale of the monopolar to bipolar transition.

### Analysis of bipolar spindle formation and spindle stability

Pole finding and tracking was implemented in a custom GUI in Matlab using previously implemented algorithms(Nazarova et al., 2013; Shulist et al., 2017). Briefly individual yeast cells were segmented using a seeded watershed algorithm, then SPs were fit to 3D gaussians and tracks were made using Hungarian based particle linking. The dual cameras were first aligned using TetraSpeck microspheres, and then chromatic shift and camera shift was measured and corrected for in the analysis.

Bipolar spindle transition velocities were computed by determining the local maxima/minima of the acceleration curves of the spindle lengths to determine start and end points of the transitions and then computing spindle formation velocity. Spindle fluctuations (Shulist et al., 2017) are computed as the standard deviation of the instantaneous spindle lengths post bipolar spindle formation as determined by the end point of the transition. Spindle fluctuations are normalized by dividing their fluctuations by their mean spindle lengths over the time course. Nascent and metaphase spindle elongation were computed by first meaning instantaneous spindle lengths with a 3-point sliding window and then using a linear regression on the data. The slope of the linear fit is used as the mean velocity of the spindle over the time course.

Spindle length equilibrium was visualized using relative probability 30 bin histograms. The probabilities where then fit to a double gaussian distribution to obtain the equilibrium distributions for the monopolar and nascent bipolar states. The cumulative distribution was computed as the length of the midpoint of the bipolar spindle formation as determined by the start and end points for the velocities.

### Analysis of delay between START and bipolar spindle formation using Whi5-GFP

Whi5-GFP nuclear export was used to identify START in G1 (unbudded) cells (Skotheim et al., 2008). Spc42-CFP was used to detect the spindle poles. Image z-stacks are max projected in FIJI (provide fiji website). START was determined by visual inspection as the first time point where the nuclear Whi5-GFP fluorescence signal was indistinguishable from that found in the cytoplasm. We determined that point to be START, as it is the completion of Whi5 export from the nucleus to the cytoplasm. The time delay until bipolar spindle formation was determined as the time elapsed between START and the first timepoint where two distinct spindle poles foci could be resolved. Estimation plots were calculated as a shared control estimation plot (Gardner and Altman, 1986) which presents the mean differences between a single WT control and Kinesin 5 mutant strains. 95% confidence interval of the mean difference between each dataset were computed and shown in relation to the WT delay.

### Electron tomography of monopolar spindles

Electron tomography was performed in collaboration with Eileen O’Toole and Mark Winey at the University of Colorado, Boulder. Unbudded (G1) cells were enriched using centrifugal elutriation on a Beckman Coulter Avanti system. Briefly, 200 mL yeast cultures were grown in YPD to an OD600 of 0.5. This culture was concentrated and injected into the centrifugal elutriator. Cells were separated using a flow rate of seven millilitres per minute and a centrifuge rotor speed of 1100 RPM. Fractions were collected every ten minutes and checked under the microscope for enrichment of unbudded cells. Fractions enriched for unbudded cells were then used to make resin-embedded tomography sections at the FEMR (McGill University). Cells from enriched fractions were collected by vacuum filtration and scraped into a 0.1 mm brass planchette. Cells were then vitrified using a Leica Microsystems EM PACT2 High Pressure Freezer System. Afterwards, cells were removed from the planchette and underwent freeze-substitution using a solution of 0.25% glutaraldehyde/0.1% uranyl acetate in 100% acetone(O’Toole et al., 2002). Cells were then embedded in Lowicryl HM20, and polymerized with UV radiation at −45°C using a Leica Microsystems EM AFS2. Resin-embedded cells were cut into serial sections, collected onto slot grids, stained at the University of Colorado, Boulder and tomography performed as previously described(O’Toole et al., 2002). Monopolar spindle models were reconstructed from tomographs using the IMOD software package(Kremer et al., 1996; Mastronarde, 1997). MT pairing analysis was performed using MatLab (MathWorks, Natick, MA). Pairwise distances between MT contour lengths were computed and overlaps defined as regions separated by ≤ 45nm over a length of ≥10nm as defined in previous studies(Nazarova et al., 2013). Overlap angles were computed from vectors formed by the start and end of the overlap region of each MT pair in order to account for bent MTs.

## 2. Supplemental tables

**Table S1.** Parameters for interactions between paired microtubules within monopolar spindles

|  | **monopolar state**<br>(n=4) | **bipolar state (800 nm)**<br>(n=1) |
| --- | --- | --- |
| **overlap number**<br>(n= MT pairs) | 48.8 ± 14<br>(n = 202) | 48 |
| **mean overlap length (nm)**<br>(n= MT pairs) | 57.7 ± 23.7<br>(n = 202) | 124.1 ± 104 |
| **mean total overlap (μm)**<br>(per spindle) | 2.82 ± 0.85 | 5.99 |
| **mean overlap angle (°)**<br>(n= MT pairs) | 83.0 ± 28.4<br>(n = 202) | 162.3 ± 27.6<br>(n = 55) |

**Table S2.**
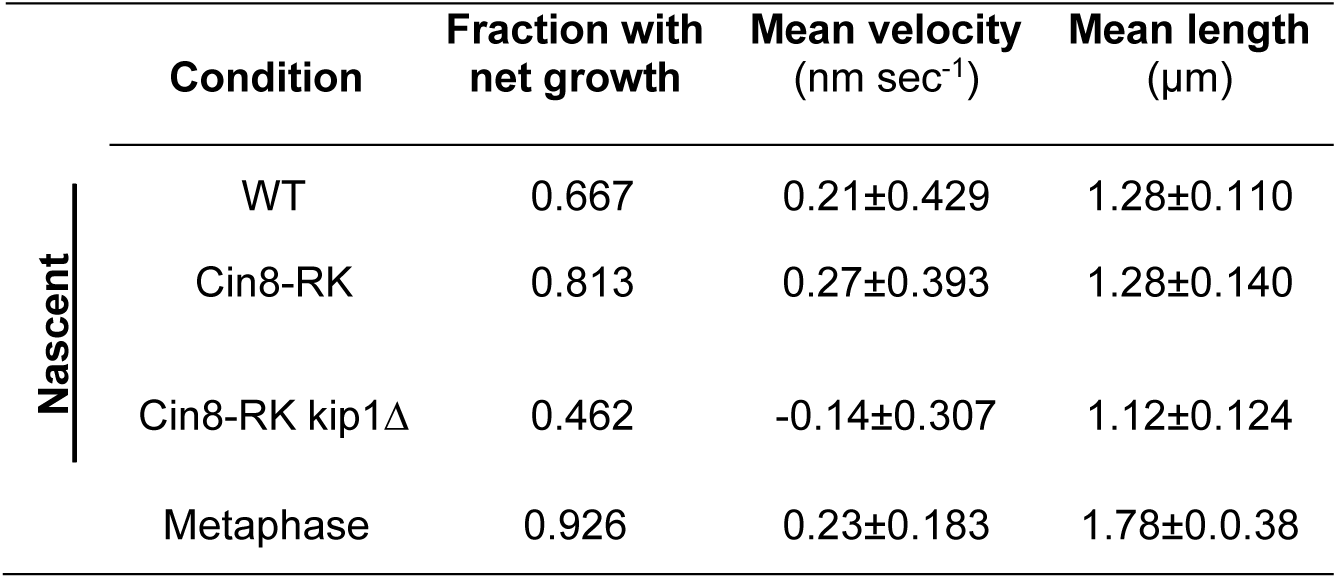
Growth parameters for nascent and metaphase spindles

|  | **Condition** | **Fraction with**<br>**net growth** | **Mean velocity**<br>(nm sec<sup>-1</sup>) | **Mean length**<br>(μm) |
| --- | --- | --- | --- | --- |
| **Nascent** | WT | 0.667 | 0.21±0.429 | 1.28±0.110 |
|  | Cin8-RK | 0.813 | 0.27±0.393 | 1.28±0.140 |
|  | Cin8-RK kip1Δ | 0.462 | -0.14±0.307 | 1.12±0.124 |
|  | Metaphase | 0.926 | 0.23±0.183 | 1.78±0.038 |

## 3. Supplemental Figures

**Fig. S1.**
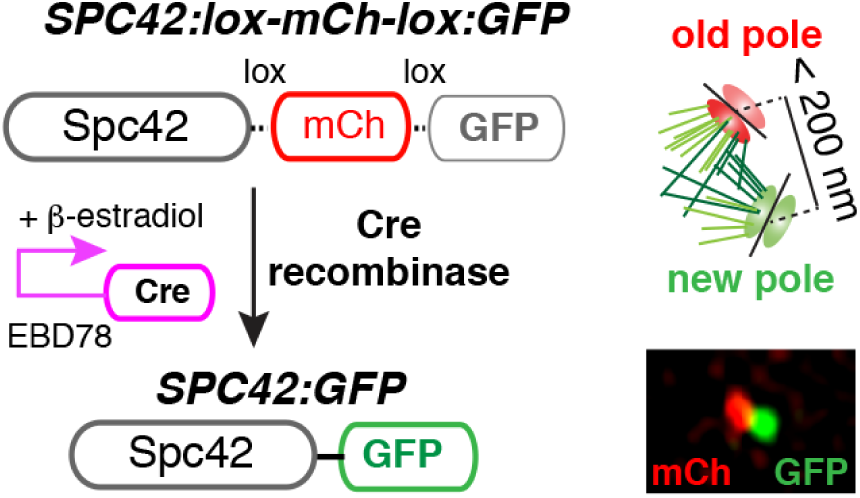
Fluorophore switching assay to resolve monopolar spindles. Cre-Lox recombinase system that recombines out an in-frame mCherry tag and leaves an in-frame EGFP tag. As a result, monopolar spindles have an old pole labelled with Spc42-mCherry (Spc42-MCh) and a new pole labelled with Spc42-eGFP, each tracked as single point-like objects during spindle formation.

**Fig. S2.**
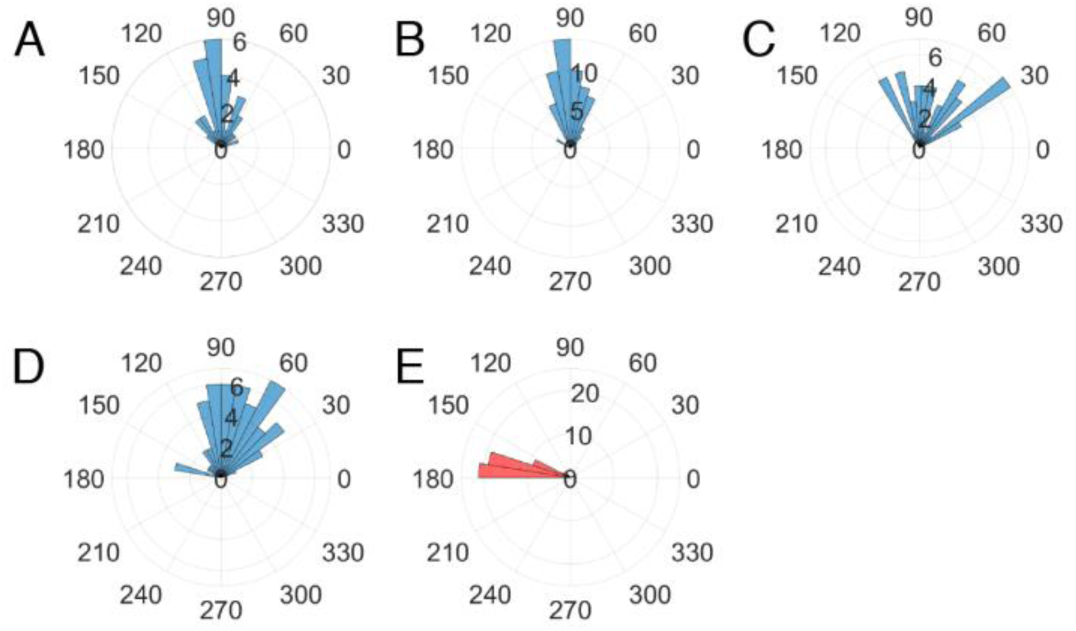
Microtubule pairing angle analysis. (A) Pre-sep1-4 and Post-sep1 from left to right. The angle formed between paired microtubules are plotted in radar plots.

**Fig. S3.**
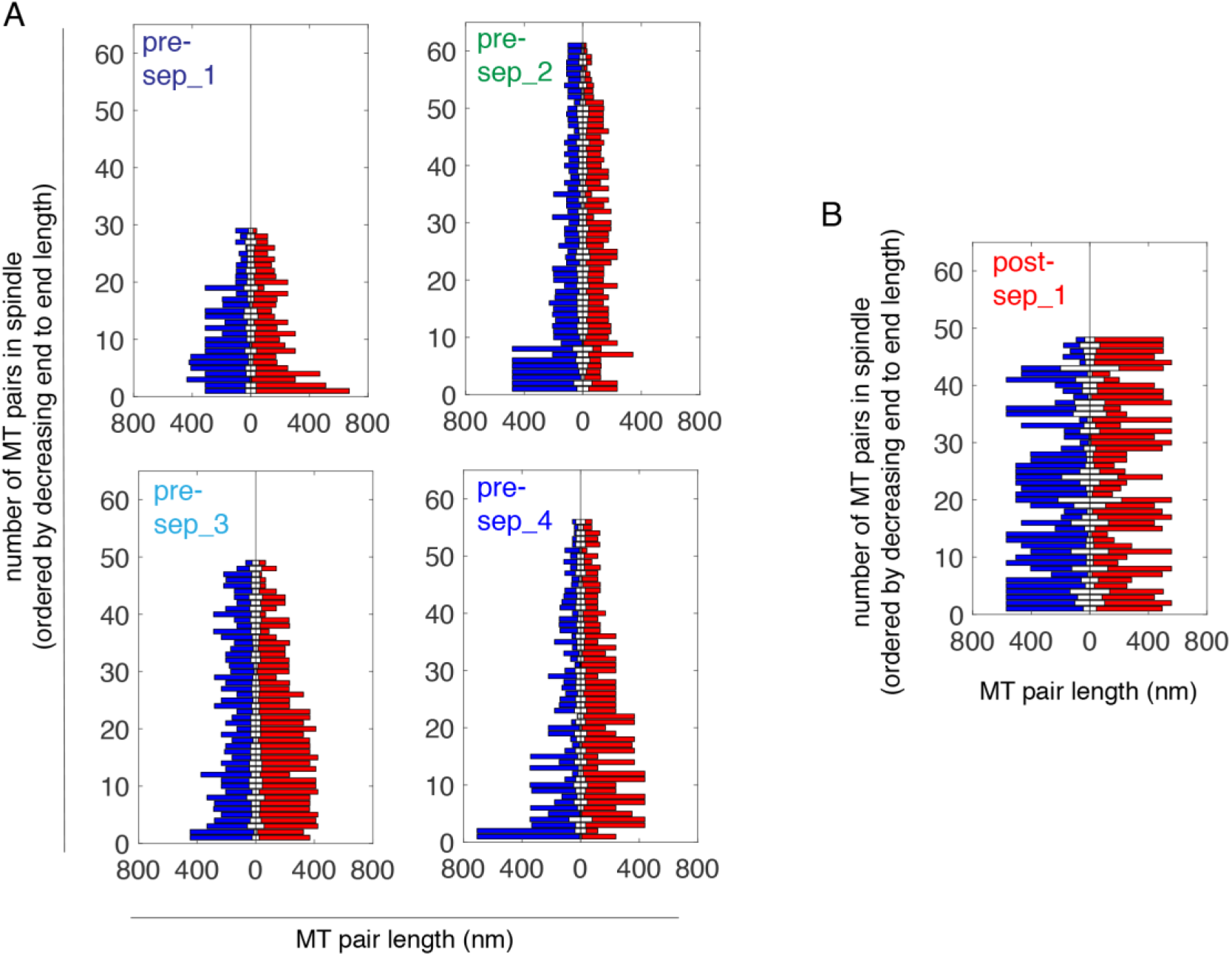
Microtubule overlap analysis. (A) Microtubule length and overlap in four monopolar spindles. Paired Microtubules emanating from the same spindle poles are represented in either blue or red. Regions of microtubule overlap are coloured in white. Microtubule pairs are ordered in ascending combined lengths. (B) Microtubule length and overlap in one short (800 nm) bipolar spindle. Paired Microtubules emanating from the same spindle poles are represented in either blue or red. Regions of microtubule overlap are coloured in white. Microtubule pairs are ordered in ascending combined lengths.

**Fig. S4.**
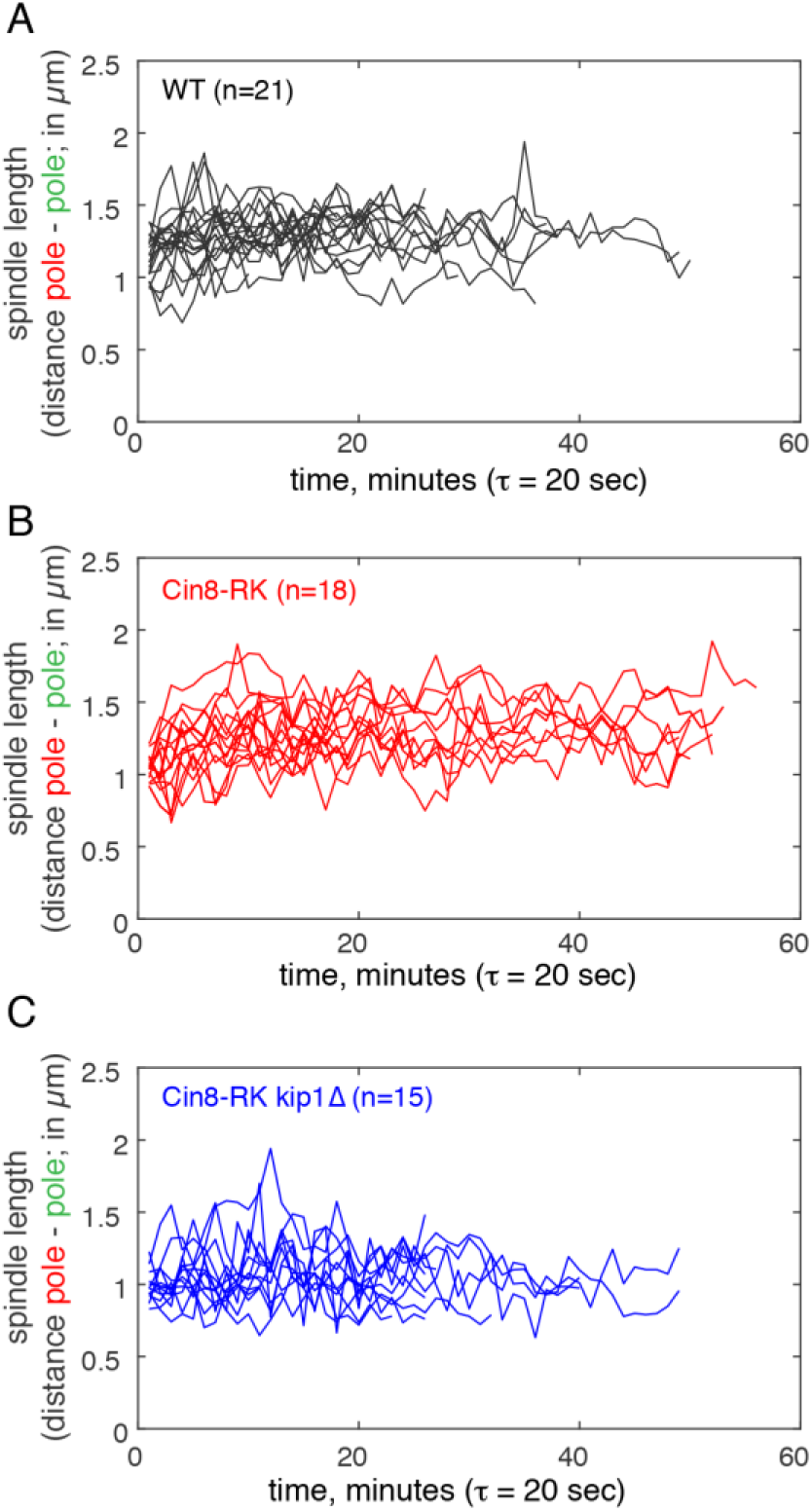
Length dynamics of nascent bipolar spindles. (A) Post-transition traces of instantaneous length of nascent bipolar spindles in WT cells. (B) Post-transition traces of instantaneous length of nascent bipolar spindles in Cin8-RK cells. (C) Post-transition traces of instantaneous length of nascent bipolar spindles in Cin8-RK kip1Δ cells.

## 4. Videos

Video 1. Time-lapse of bipolar spindle formation in live budding yeast cells

Spindle pole bodies are tagged using fluoro-switching. Spc42-mCherry-GFP. Images are taken every 20s. Inset shows the brightfield cell taken every 200s.

Videos 2-5: Tomographic reconstructions of four monopolar spindles with microtubules Microtubules emanating from the same spindle pole are coloured in either red or blue.

Video 6: Tomographic reconstruction of a short bipolar spindle with microtubules Microtubules emanating from the same spindle pole are coloured in either red or blue. (Post-sep1)

Video 7: Time lapse of the delay between START and bipolar spindle formation. Whi5-GFP nuclear exit marks the START point in the cell cycle after which the delay until bipolar spindle formation labelled as Spc42-CFP can be quantified.

